# Disentangling adaptation from drift in bottlenecked and reintroduced populations of Alpine ibex

**DOI:** 10.1101/2021.01.26.428274

**Authors:** D.M. Leigh, H.E.L. Lischer, F. Guillaume, C. Grossen, T. Günther

## Abstract

Identifying local adaptation in bottlenecked species is essential for effective conservation management. Selection detection methods are often applied to bottlenecked species and have an important role in species management plans, assessments of the species’ adaptive capacity, and looking for responses to major threats like climate change. Yet, the allele frequency changes driven by selection and exploited in selection detection methods, are similar to those caused by the strong neutral genetic drift expected during a bottleneck. Consequently, it is often unclear what accuracy selection detection methods may offer within bottlenecked populations. In this study, we used simulations to explore if signals of selection could be confidently distinguished from genetic drift across 23 bottlenecked and reintroduced populations of Alpine ibex (*Capra ibex*). We used the meticulously recorded demographic history of the Alpine ibex to generate a comprehensive simulated SNP data. The simulated SNPs were then used to benchmark the confidence we could place in putative outliers identified through selection scans on empirical Alpine ibex SNP data. Within the simulated dataset, the false positive rates were high for all selection detection methods but fell substantially when two or more selection detection methods were combined. However, the true positive rates were consistently low and became essentially negligible after this increased stringency. Despite the detection of many putative outlier loci in the empirical Alpine ibex RADseq data, none met the threshold needed to distinguish them from genetic drift-driven false positives. Unfortunately, the low true positive rate also creates a paradox, by preventing the exclusion of recent local adaptation within the Alpine ibex.

## Introduction

Identification of recent responses to selection, or local adaptation, is of great interest to evolutionary and conservation biologists. Insights gained from recent selective changes can facilitate our understanding of evolutionary processes (Whitlock and Lotterhos, 2015a). For conservation biologists, insights into local adaptation also have a more applied or practical importance. Characterizing within species adaptive differences is often necessary for species management plans (e.g. Robertson *et al.,* 2014), and optimizing source population choice for translocations or reintroductions (Flanagan *et al.,* 2017). Characterizing adaptive processes may also offer insight into long-term extinction risk, particularly if a population or species is no longer able to respond to selection (Frankham *et al.,* 2010). Within reintroduced populations specifically, the sudden environmental change experienced when founder individuals are released in new locations may fuel rapid adaptive change (e.g. Stockwell *et al.,* 2003; Reznick *et al.,* 2004). Understanding of which is important if future- potentially disruptive- translocations are planned. This new conservation ethos where evolutionary processes are considered in species management, is known as “evolutionary” or “adaptive” conservation management (Hoffmann *et al.*, 2015). The long-term success of evolutionary conservation management requires accurate assessments of the evolutionary processes in bottlenecked populations and thus, an understanding of the analytical constraints non-equilibrium populations can face.

The current ease in obtaining genome-wide SNP data has driven a renaissance of studies scanning for selection at the genomic level in wild populations (e.g. *Gasterosteus aculeatus*, Hohenlohe *et al.,* 2010; *Peromyscus maniculatus*, Linnen *et al.,* 2013; *Sarcophilus harrisii,* Epstein *et al.,* 2016; *Oncorhynchus clarkii henshawi,* Amisch *et al.,* 2019). *Fst*-based selection detection methods are widely used to detect recent intra-species selective responses by scanning for unusually high values of *Fst* (“outlier” loci), which are assumed to be driven directly or indirectly (i.e. hitchhiking) by positive selection (Lewontin and Krakauer, 1973; Fay and Wu, 2000). Popularity of these methods has fueled analytical extensions that identify selective responses using environmental clines (Coop *et al.,* 2010; De Mita *et al.,* 2013). Referred to as genetic-environment association analyses or “GEA” analyses, these methods pinpoint alleles that display repeated associations with an environmental variable due to local adaptation (Lotterhos and Whitlock, 2015; Hoban *et al.,* 2016). The degree to which currently available selection detection methods successfully accommodate unusual, or more complex demographic histories, is still being tested. This information is essential to ensure accuracy because small demographic assumption violations can fuel elevated rates of false signals of selection, where neutral loci are falsely identified as outliers. This can arise, for example, from unaccounted variance in the distribution of *Fst* due to shared history and relatedness of populations (Robertson, 1975a; Robertson, 1975b; Excoffier *et al.,* 2009). Recent population bottlenecks and reintroductions pose a new challenge for selection detection, because they are associated with very complex patterns of high inter-population relatedness that may violate model assumptions and exacerbate false positive rates (Frankham *et al.,* 2010). Furthermore, the random allele frequency changes caused by the strong genetic drift inherent in a bottleneck can lead to large allele-frequency differences between populations (Kimura 1955a; Kimura 1955b). Genetic drift can therefore create outlier-like loci that can easily be mistaken as loci under selection and will increase the false positive rate of selection detection methods in bottlenecked populations (Lotterhos and Whitlock, 2014; Klopfstein *et al.,* 2006; Nielsen *et al.,* 2007; Foll and Gaggiotti, 2008; Hofer *et al.,* 2009). Such false signals have previously hampered selection scans in bottlenecked species, including humans (Sabeti *et al.,* 2006).

Examination of selection detection accuracy in bottlenecked populations is limited. Foll and Gaggiotti, (2008) examined the effects of including a subset of populations that are bottlenecked in a selection detection analysis. It was recommended to remove bottlenecked populations due to the increase in false positives this caused (Foll and Gaggiotti, 2008). The effects of historical bottlenecks (thousands of generations prior) were also examined using simulated populations of *Peromyscus spp.* (Poh *et al.,* 2014) and *Haemorhous mexicanus* (Shultz *et al.,* 2016), where the false positive rate often exceeded selection detection power. Nevertheless, selection detection analyses have since been applied to bottlenecked populations (e.g. Pilot *et al.,* 2014; Funk *et al.,* 2016; Amish *et al.*, 2019), and will likely continue to be applied, because of the conservation management need to identify intra-species adaptive differences. It is therefore essential that we expand our exploration of bottleneck effects on selection detection accuracy.

The Alpine ibex (*Capra ibex)* is a recently bottlenecked and reintroduced species with a demographic history that is virtually unparalleled in recorded detail (Biebach and Keller, 2009). In this study, we utilized these population records to create a comprehensive simulated SNP data set through individual-based forward simulations. We then examined the performance of different selection detection methods by quantifying both the observed true and false positive rates and the composition of outlier loci. This information was coupled with selection scans on an empirical Alpine ibex restriction site associated DNA sequencing (RADseq) data set, and used to guide the confidence we could place in any outliers detected in these reintroduced populations. This provided insight into the accuracy, or rather lack-there-of, expected within species with complex histories of bottlenecks and reintroductions. The detection thresholds and methods outlined here can be used as a guideline to help avoid false positive loci in other species with similar histories.

## Materials and methods

### Alpine ibex demographic history

Alpine ibex underwent a prolonged decline starting in the 16^th^ century due to overhunting. Only a single population of an estimated 100 individuals survived this crash in the Gran Paradiso region of Northern Italy. Royal protection in the 19^th^ century enabled the population to grow to 3000-5000 individuals. Reintroductions of Alpine ibex from the Gran Paradiso region into Switzerland began in 1906. Detailed demographic records were kept as part of the reintroduction program in Switzerland. Information that was recorded included the origin of founder individuals (often coming from previously reintroduced populations, Figure 1), the number and gender of founders, and the year an individual was moved. In addition, annual census records of the number of animals alive in spring were collected for many reintroduced populations (Stuwe and Grodinsky, 1987; Stuwe and Neivergelt, 1991; Biebach and Keller, 2009). This reintroduction program was very successful, to date more than 17 thousand Alpine ibex are present in the Swiss Alps (Shackleton and Group ISCI, 1997; BAFU, 2015; Brambilla *et al.* 2020). The focal populations used in this study are shown in Figure 1.

**Figure 1:**
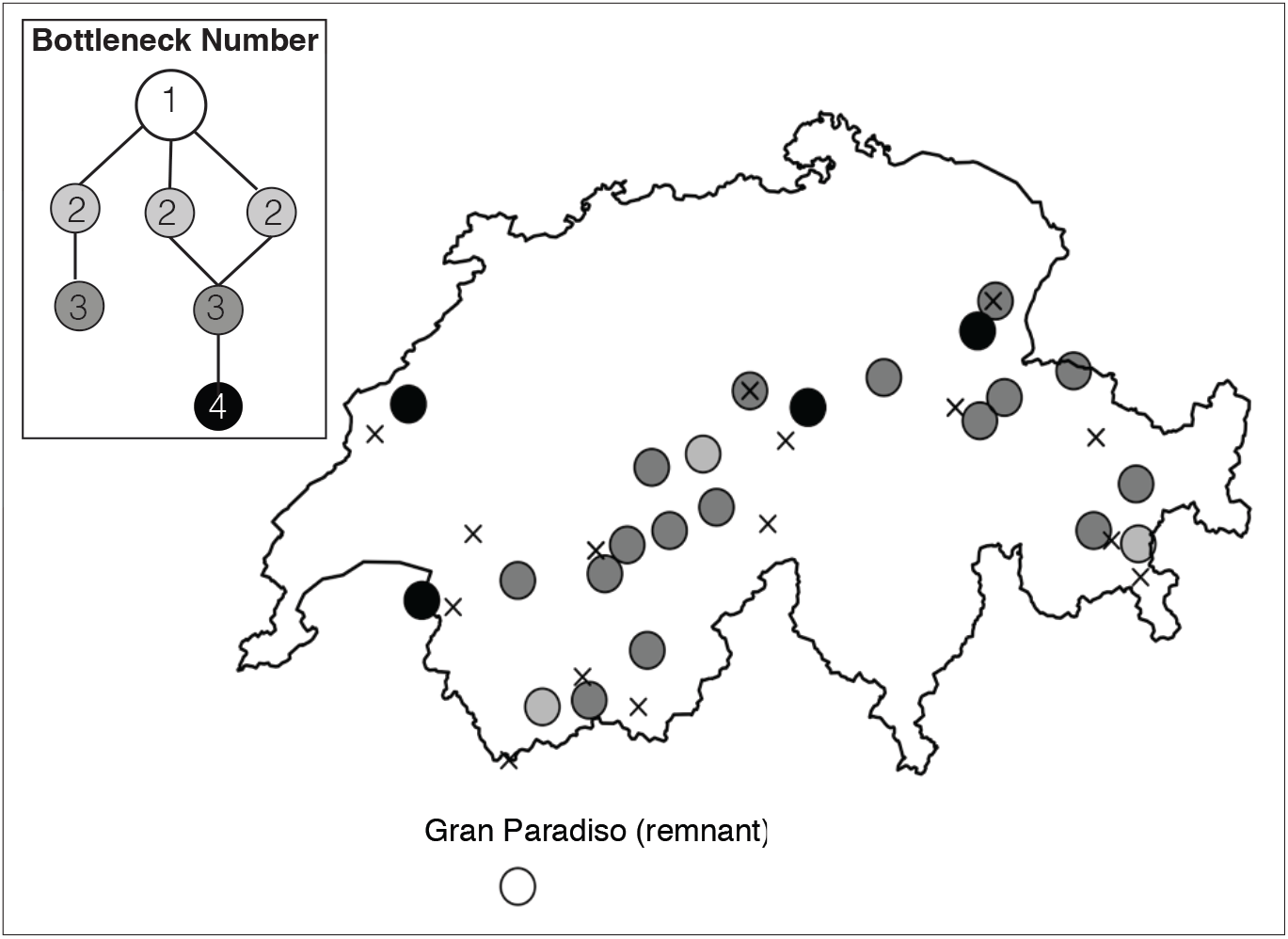
The 23 Alpine ibex focal populations and a simplified representation of the reintroduction history in Switzerland equating to the effective bottleneck number each population experienced (top left panel). All Swiss populations descend from the Gran Paradiso national park in Northern Italy (open circle), which is included in the figure but was excluded from the selection detection analysis. Reintroductions in Switzerland often used founder individuals from previously established reintroduced populations. As a result, many populations have experienced several serial bottlenecks. Within this figure, each circle represents a Swiss Alpine ibex focal population and the circle’s shading indicates the number of bottlenecks each population experienced. Marked by a cross are the weather stations used to estimate the local environment experienced by each population.

### RAD sequencing

To apply selection detection methods to an empirical data set from a bottlenecked species, we used the published RADseq data set from Leigh *et al.,* (2018) and Grossen *et al.,* (2017). This consists of 304 Alpine ibex from 23 reintroduced populations (Figure 1). We used only variants called by GATK (Poplin *et al.,* 2017; see Leigh *et al.,* 2018 for a discussion of variant caller effects). After SNP filtering (described in section S3) a sample of 213 individuals remained. For selection detection all singletons were removed and SNPs within 1kb were randomly thinned using vcftools (vcftools; Danecek *et al.,* 2011), which resulted in a final data set of 12695 SNPs. After exclusion of individuals from the Gran Paradiso, inclusion of which potentially violates selection detection analysis relatedness assumptions (Günther and Coop, 2013), 5225 SNPs were suitable for the selection detection analyses.

### Simulating the Alpine ibex history

Simulated SNP data sets were generated using forward time simulations in Nemo (version 2.3.51; Guillaume and Rougemont, 2006) and used to assess the expected accuracy if each selection detection methods when applied to bottlenecked and reintroduced species. Details of the simulations can be found in the supplementary material (S1). Briefly, in each simulation all 23 populations sampled for RADseq were simulated. In order to accurately simulate these populations, an additional three populations that were founder sources for the focal populations were also simulated (see panel in Figure 1). Therefore, 26 populations were simulated in total. The reintroduction history and population sizes were informed by detailed records and census data. Ten replicate simulations of the Alpine ibex reintroduction history were conducted for each of three genetic architectures: 1) neutral SNPs only, 2) 30 loci under selection, and 3) 120 loci under selection. The loci under selection were di-allelic QTL contributing additively to a quantitative trait. In all architectures, each individual had 30 chromosomes (linkage groups) of 10M (Morgan) each with 60 thousand neutral loci. In the two architectures with selection the 30 or 120 QTL were equally spread among the neutral loci. The recombination rate was 5×10^−4^ between adjacent neutral SNPs. The QTL were set either at the center of each chromosome (30 QTL) or four QTL were positioned 3.33M apart and 0.5cM from the start on each chromosome (120QTL). This ensured several thousand SNPs were polymorphic after the bottleneck and generated the same chromosome number and a similar level of linkage disequilibrium to that in the RADseq data set as evaluated by the r^2^ values between final polymorphic SNPs in vcftools.

In each simulation, neutral loci and loci under selection were allowed to reach mutation-selection-drift equilibrium during a “burn-in” of 10 thousand generations in a single population that represented the Gran Paradiso population. After this time, a bottleneck was applied. We simulated phenotypic selection on the quantitative trait with a Gaussian fitness surface where the trait optimum value varies among populations depending on an environmental variable (snow cover). The trait optimum value during the burn-in was held at zero (in the ‘Gran Paradiso’ reference population) to maintain alleles of both negative and positive effect. To generate post-reintroduction selection across the 30 or 120 QTL, the trait optimum in reintroduced populations was varied to either zero, −2 or +2. Values reflected observed real world snow conditions relative to the Gran Paradiso, for example those with a higher average snow depth had an value of +2 and those with a lower average snow depth had a value of −2. Snow conditions were chosen as they are a strong candidate real-world selection pressure, specifically they have previously been shown to affect Alpine ibex population dynamics and vary dramatically across sites (detailed in S1 and S2) (Jacobsen *et al.,* 2004; Grøtan *et al.,* 2008).

The strength of selection at each locus was determined by the size of its contribution to the trait. For the architecture where 30 diploid loci were under selection: six loci had large contributions to each trait (allelic value, *a* = ±0.1), and 24 were divided equally into 4 categories of lesser effect (*a* = ±0.08, ±0.04, ±0.02, ±0.01). A maximum trait value of ±3 was therefore achievable. For the architecture where 120 loci were under selection, the division of loci remained identical except for the loci of smallest effect. Specifically, 96 loci were of minor effect (±0.01) and 24 were equally divided amongst the remaining allelic values (±0.1, ±0.08, ±0.04, ±0.02, 6 of each value in total). A maximum trait value of ±4.8 was achievable. Selection coefficients (*s*) equaled 0.027 (*a* = ±0.1), 0.022 (*a* = ±0.08), 0.012 (*a* = ±0.04), 0.007 (*a* = ±0.02) and 0.004 (*a* = ±0.01) in both architectures. This was calculated according to Bürger (2000) using the phenotypic variance (Vp) of 0.047 (120 loci under selection) or 0.035 (30 loci under selection), as well as a selection variance (ω^2^) of 7.5. This generated two biologically realistic trait architectures and realistic strengths of selection.

The simulated genotypes from the final generation were used to evaluate the expected accuracy of different selection detection methods, and only polymorphic SNPs were included in the simulated data from this time point. To mimic the available RADseq data, 10 simulated individuals were randomly chosen from each of the 23 populations that were sequenced with RADseq, 6000 polymorphic loci were the taken for each individual including all polymorphic selected loci and a subset of randomly selected neutral loci. 20% of genotypes were randomly set to “missing” due to missing data in the RADseq genotypes and singletons were removed (vcftools; Danecek *et al.,* 2011). PGDspider (version: 2.0.9.2; Lischer and Excoffier, 2012) and custom scripts were used to convert Nemo output into input for the selection analyses.

### Screens for signals of positive selection

Selection detection analyses were conducted for both the empirical Alpine ibex RADseq data and simulated data sets. This enables us to quantify the confidence we could place in any empirical outliers. To detect signatures of selection, Bayenv 2.0 (Günther and Coop, 2013), Baypass 2.1 (Gautier, 2015a), and OutFLANK (Whitlock and Lotterhos, 2015a) were used (following Leigh *et al.,* 2018). These three programs were chosen as they have been shown to have high accuracy in species with complex patterns of population relatedness (Günther and Coop, 2013; Lotterhos and Whitlock, 2014; Gautier, 2015a; Whitlock and Lotterhos, 2015a). Bayenv 2.0 and Baypass2.1 utilize a modified *Fst*-like statistic called X^T^X that is corrected for shared population history (Günther and Coop, 2013; Gautier, 2015a). Outflank utilizes an *Fst* statistic called *F’st*, a metric based on Wright’s *Fst* statistic without corrections for a finite sample size (Whitlock and Lotterhos, 2015a). These three methods are hereafter referred to as *Fst*-like approaches. Bayenv 2.0 and Baypass2.1 also detect selection using GEA selection scans (as in Hoban *et al.,* 2016).

Selection detection program conditions are detailed in Leigh *et al.,* (2018). Briefly, the estimation of covariance matrix and subsequence selection scan in Bayenv 2.0 was run independently three times with 2 x10^5^ Markov-Chain-Monte-Carlo (MCMC) iterations (Blair *et al*., 2014). SNPs were considered putatively under selection for the GEA method, if the Bayes factor (BF) value exceeded 3 and the Spearman’s rho value was in the top and bottom 2.5% of all SNPs across the three runs. This threshold was chosen because it suggests high support for a SNP being under selection and that the trend is not due to a single outlier population (Nadeau *et al.,* 2016; Günther and Coop, 2013). The *Fst*-like approach SNPs had to have X^T^X value among the top 100 ranking SNPs across all three runs (Günther and Coop, 2013).

Baypass2.1 was run three times for each data set with 20 pilot runs of 1000 MCMC iterations and 5000 MCMC iterations for the “burn-in” (default conditions). For the GEA analysis we used the Auxillary model and consider a loci to be under selection when it had a 10 x log10 Bayes factor (db) greater than 4.7 for all three replicates (Gautier, 2015a). This value is equivalent to the threshold of a BF of 3 used in Bayenv 2.0. For the *Fst*-like approach, X^T^X outliers were determined following the best-practice tutorial accompanying Baypass2.1 (Gautier, 2015b). This uses trained-simulations to find the 99% threshold for X^T^X values for each dataset, outliers were those loci in the top 1% for all three Baypass runs (Gautier, 2015b).

In OutFLANK, outlier SNPs were identified following the best practice tutorial (default settings, Whitlock and Lotterhos, 2015b). To be considered an outlier, a SNP had to have a Q-value of less than 0.05 (Storey and Tibshirani, 2003; Whitlock and Lotterhos, 2015a), as well as a heterozygosity of greater than 10% (Whitlock and Lotterhos, 2015b).

Loci identified across multiple programs as outliers were also compared. Loci identified as outliers across two programs were called “double positives” those found by all three programs were called “triple positives.” To account for the different signals the *Fst*-like and GEA approaches look for, the outliers identified by the two methods in Bayenv 2.0 and Baypass2.1 were not combined into a single set. Thus we had double and triple positive *Fst*-like outliers, and double positive GEA outliers. For the triple positive GEA outliers, the GEA outliers from Bayenv 2.0 and Baypass2.1 were overlapped with the *Fst-*like outliers from OutFLANK because OutFLANK does not us a GEA approach.

All environmental data used in the GEA analyses were obtained from MeteoSwiss (Switzerland). For each population, data from the closest meteorological station available (Figure 1, Section S1 and S2) were used to obtain averages since a population was founded, or since records began. The environmental variables in the analyses were divided across winter and summer and included air temperature (°C), daily precipitation (mm), and snow depth measures (cm). Further details are available in the supplementary material (section S1 and S2). Since the simulations were intended to mimic real Alpine ibex populations, the corresponding weather data were included as environmental covariates in the Bayenv 2.0 and Baypass2.1 analyses of the simulated data. In addition, each simulated population’s true simulated environmental optimum was also included as an environmental covariate in the analysis of the simulated data (Table S1).

### Evaluating method accuracy with simulations

The simulated genotype data was used to estimate the true or false negative and positive rates. When examining loci flagged as putatively under selection, a true positive was considered to be a simulated locus under selection that was correctly identified as being under selection. A false positive was considered to be a simulated neutral locus that was wrongly identified as being under selection. The proportion of all loci identified by a test as under selection that were true positives, hence indeed under selection (the true discovery rate), was used as a metric of the accuracy and reliability of selection detection. To place the results in the context of other simulation studies, the true positive rate, false positive rate, the false discovery rate, and false negative rate, were also calculated. All metrics are defined in Table 1 for ease of reference. All values displayed are the averages across 10 simulated datasets for each genetic architecture and are relative only to the number of polymorphic QTL loci and neutral loci in the final SNP set.

**Table 1:** Definitions of each metric used to assess a selection detection method’s accuracy with the simulated data.

| Accuracy metric | Definition |
| --- | --- |
| *True discovery rate* | The proportion of all simulated loci identified as outliers that were actually under selection (i.e. QTL loci). |
| *True positive rate* | The proportion of loci under selection (i.e. QTL loci) correctly identified as an outlier. |
| *False positive rate* | The number of neutral loci incorrectly identified as under selection (false positive) divided by the number of retained polymorphic neutral SNPs. |
| *False discovery rate* | The proportion of outlier SNPs that were false positives (i.e. simulated neutral loci)(Lotterhos and Whitlock, 2014) |
| *False negative rate* | The proportion of polymorphic QTLs that were not identified as outliers (and thus not identified as under selection). |

## Results

In this study, we generated empirical RADseq and simulated SNP data for the Alpine ibex. Bayenv 2.0, Baypass2.1, and OutFLANK were then used to identify loci putatively under selection in these datasets. The simulated data provided an estimate of the selection detection accuracy of these three popular tools in the empirical Alpine ibex dataset. Low true discovery rates were identified for all selection detection methods (detailed below), preventing us from confidently distinguishing selection from false positive outliers in the Alpine ibex RADseq data.

### Alpine ibex RADseq data and signals of selection

Each selection detection software identified outliers in the Alpine ibex RADseq data set. Between 172 to 2 loci were found to be putatively under selection by the different selection detection methods (Figure 2A). However, only 14 loci were identified as double positives and no locus exceeded the triple positive threshold. The highest number of double positive loci was found by the Bayenv Baypass GEA overlap. The two other double positive loci were found separately in the overlap of Bayenv and Baypass *Fst*-like, as well as the Bayenv and Outflank *Fst-*like overlap. As detailed below, this is within the range of drift-driven false positives expected under all simulated genetic architectures.

**Figure 2:**
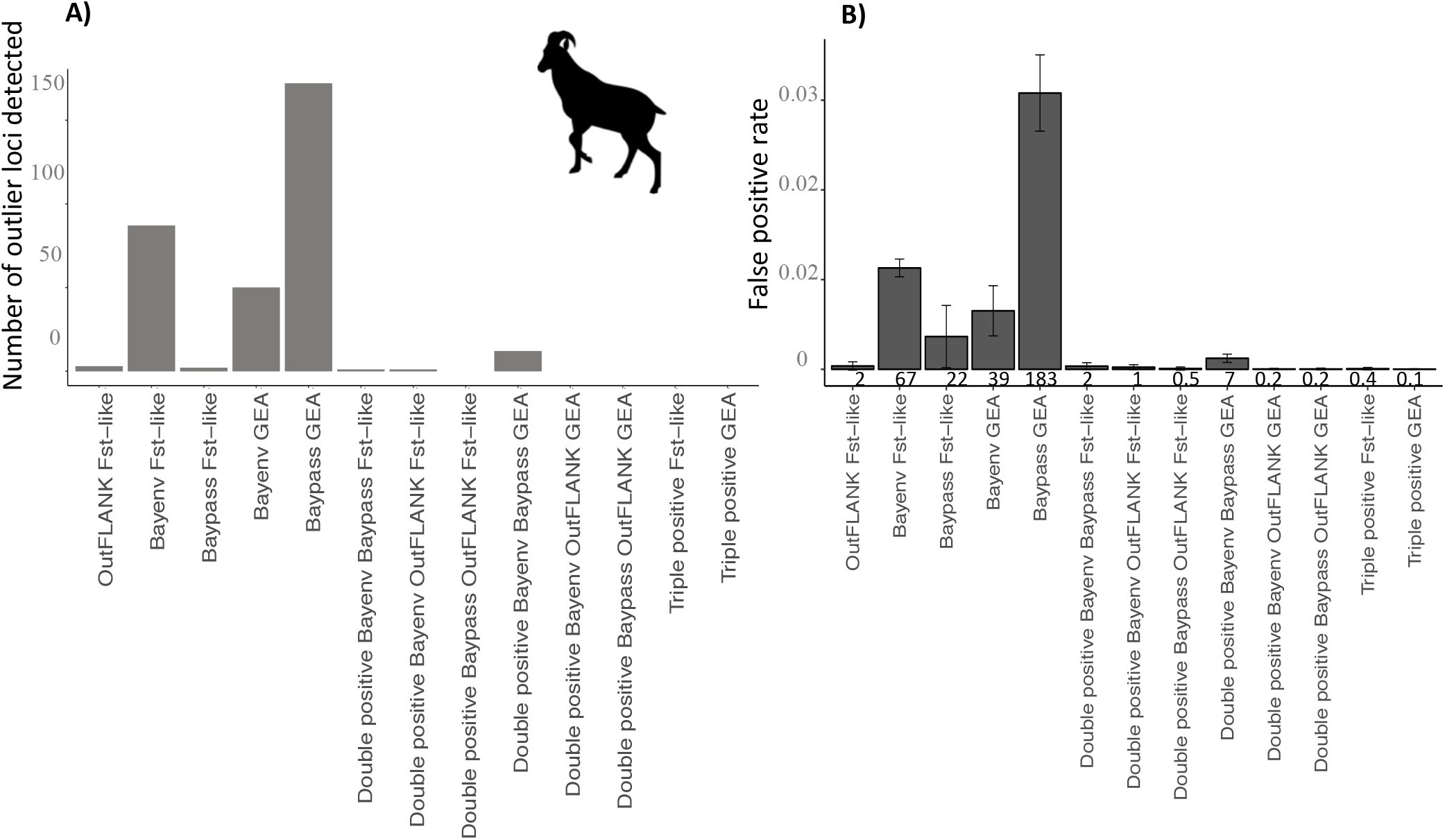
**A)** The number of empirical outliers detected by each selection detection method in the Alpine ibex RADseq SNP set. **B)** The false positive rate from the fully neutral simulations. Shown below each bar is the average number of outlier loci identified

### Evaluating expected selection detection accuracy

Analyses of simulated data revealed a very low selection detection accuracy under the Alpine ibex demography, regardless of the genetic architecture simulated. Figure 2B shows the false positive rates for the neutral only simulation and Figure 3 the true and false discovery rates (i.e. the composition of loci identified as outliers) for the simulations with loci under selection. For the two architectures with selection, the true positive rate, false positive rate and false negative rates are shown in Table 2.

**Figure 3:**
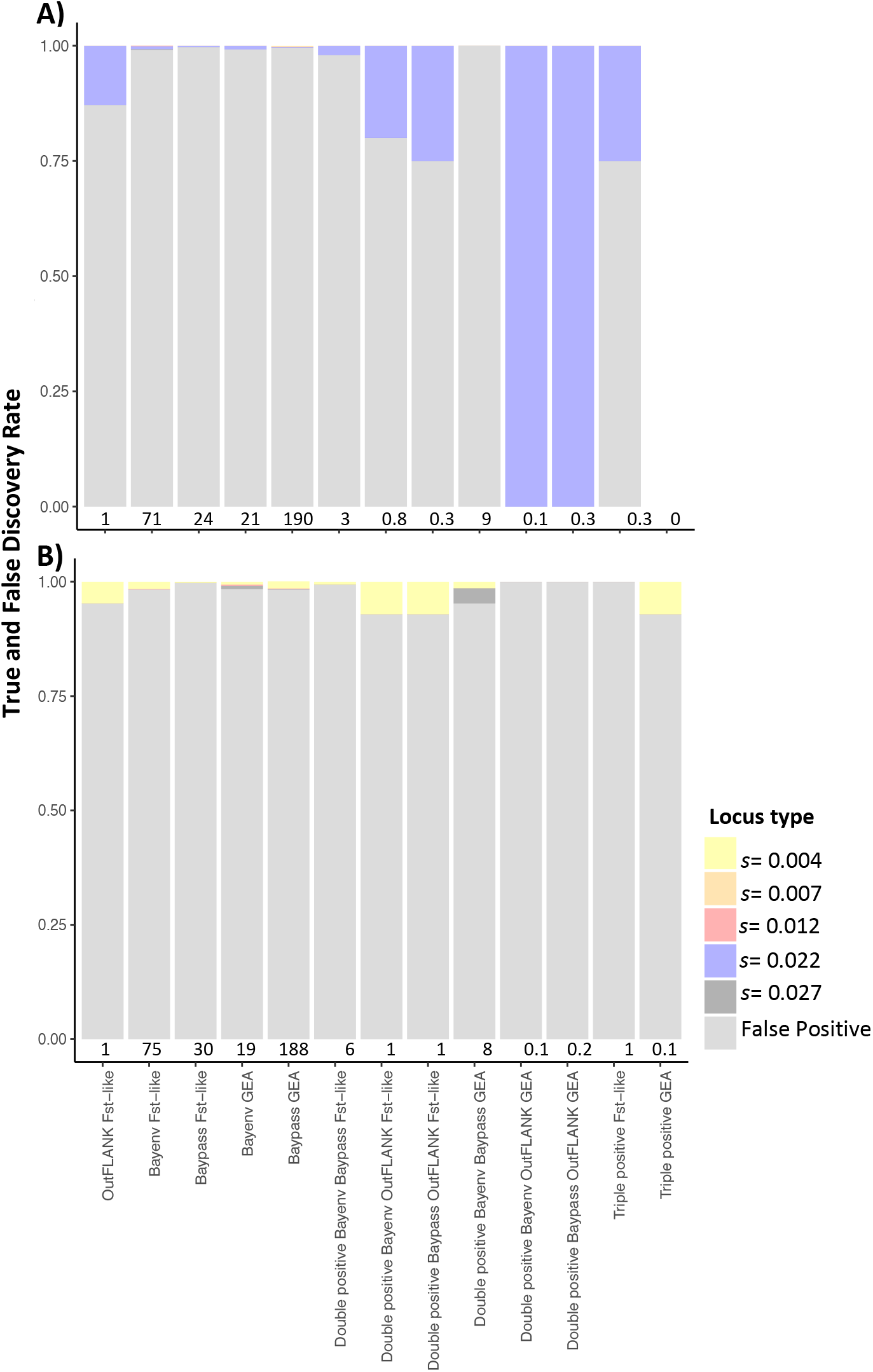
The true and false discovery rate of different selection detection methods for **A)** the architecture with 30 loci under selection and **B)** the architecture with 120 loci under selection. Each bar shows the average composition of loci identified as outliers using each selection detection method, at the bottom of the bar is the average number of outliers across 10 replicate simulations. Replicates where no loci exceeded the significance threshold were excluded from the figure.

**Table 2:** Selection detection accuracy as measured by the true and false positive rate, as well as the false negative rate. 30 or 120 signifies the number of loci under selection (QTL loci).

| Selection detection method | True positive rate |  | False positive rate |  | False negative rate |  |
| --- | --- | --- | --- | --- | --- | --- |
|  | 30 | 120 | 30 | 120 | 30 | 120 |
| Bayenv *Fst-like* | 0.026 | 0.012 | 0.012 | 0.012 | 0.974 | 0.988 |
| Baypass *Fst-like* | 0.005 | 0.001 | 0.004 | 0.005 | 0.995 | 0.999 |
| OutFLANK *Fst-like* | 0.009 | 0.001 | 0.000 | 0.001 | 0.991 | 0.999 |
| Bayenv GEA | 0.005 | 0.004 | 0.003 | 0.003 | 0.995 | 0.996 |
| Baypass GEA | 0.033 | 0.030 | 0.032 | 0.031 | 0.967 | 0.970 |
| Double positive Bayenv Baypass *Fst-like* | 0.005 | 0.001 | 0.001 | 0.001 | 0.995 | 0.999 |
| Double positive Bayenv OutFLANK *Fst-like* | 0.009 | 0.001 | 0.000 | 0.001 | 0.991 | 0.999 |
| Double positive Baypass OutFLANK *Fst-like* | 0.005 | 0.001 | 0.000 | 0.001 | 0.995 | 0.999 |
| Double positive Bayenv Baypass GEA | 0.000 | 0.003 | 0.001 | 0.001 | 1.000 | 0.997 |
| Double positive Bayenv OutFLANK GEA | 0.005 | 0.000 | 0.000 | 0.000 | 0.995 | 1.000 |
| Double positive Baypass OutFLANK GEA | 0.005 | 0.000 | 0.000 | 0.000 | 0.995 | 1.000 |
| Triple positive *Fst-like* | 0.005 | 0.001 | 0.000 | 0.001 | 0.995 | 0.999 |
| Triple positive GEA | 0.000 | 0.000 | 0.000 | 0.000 | 1.000 | 1.000 |

For all simulation types, each individual selection detection method had a high number of false positives and a striking false negative rate (Figures 3). The false positives rate did decrease considerably (<0.001) for the double and triple positive methods, but this was at the expense of the false negative rate increasing (Table 2). Greater variability in accuracy is seen for the architecture with 30 loci under selection than 120 loci under selection. Specifically, the true discovery rate does occasionally reach 1.0 (see Figure 3). However, as shown by the true positive rate and false negative rate (Table 2), this does not reflect high accuracy of these methods but stochastic chance. Virtually all simulations had no outliers exceed this threshold, but a single simulation had 1 true positive locus, leading to a mean true discovery rate of 1.

In the simulations with selection, the allelic values and hence the strength of selection experienced by each QTL locus were not equal. The loci with allelic values of 0.1 or 0.08 were under much stronger selection (*s=*0.027, 0.022) relative to those with allelic values of 0.04, 0.02 or 0.01 (*s=*0.012, 0.007, 0.004). Consequently, the signal of selection and therefore the true positive rate may be unequal across loci under selection. Table 3 shows the average allele frequency change of loci under selection, this can be considered a rough proxy for the signal of selection visible at a locus. As expected due to the strength of selection, loci under the strongest selection were often at extreme allele frequencies after the burn-in and before the bottleneck (Figure S1 and S2). Consequently, such loci were fixed more frequently over the course of our simulations and thus more likely to be excluded from selection detection analysis. Nevertheless, loci under a selection pressure of >0.022 were the most likely to be identified as outliers in the architecture with 30 loci under selection. However, those under weaker selection (0.004) were most likely to be identified as outliers in the architecture with 120 loci under selection but this was because they were by far the most common in this architecture, their abundance drives this trend.

**Table 3:** Mean absolute allele frequency change for loci under selection ± the standard error. Shown in brackets is the percentage of loci that remain polymorphic in at least one population at the end of the simulations. Values are calculated from immediately after the burn-in using the values from the simulated Gran Paradiso population, relative to the frequency across all simulated populations in final generation. Loci fixed after the burn-in were excluded from the values.

| Locus type | Average allele frequency change (percentage polymorphic) |  |
| --- | --- | --- |
|  | 30 Loci under selection | 120 Loci under selection |
| 0.01 | 0.086±0.071 (93%) | 0.087±0.079 (86%) |
| 0.02 | 0.095±0.082 (94%) | 0.093±0.078 (93%) |
| 0.04 | 0.067±0.067 (77%) | 0.088±0.078 (83%) |
| 0.08 | 0.043±0.058 (48%) | 0.045±0.060 (49%) |
| 0.1 | 0.031±0.034 (44%) | 0.016±0.022 (23%) |

## Discussion

In this study the accuracy of selection detection methods was assessed for the Alpine ibex, a species with a complex history of bottlenecks and reintroductions. We generated comprehensive simulations that followed the species’ recorded population history. Three genetic architectures were simulated: neutral loci only, 30 loci under selection, and 120 loci under selection. The simulated data revealed a low selection detection accuracy for each individual selection detection method. Improved accuracy was possible when only considering outliers identified by multiple methods, though this came at the expense of an increased false negative rate. This made it impossible to adjust our thresholds as we were either overrun with false positives, or rarely identified ongoing selection. While candidate outlier loci could be identified in the Alpine ibex RADseq data set, the simulation results indicate they cannot be confidently considered as under selection. Importantly, the low true positive rate also prevents us from confidently concluding the absence of recent adaptation in the populations, posing significant challenges for the evolutionary management of this species. Nevertheless, identifying false positive outliers and concluding two populations are separate ESUs has a number of costly consequences for conservation management. Until more accurate selection detection methods are found, the stringent approach and criteria here should be applied to other bottlenecked species to offer an indication of the confidence that we can place in outlier loci.

### Screen for selection with Alpine ibex RADseq data

In the Alpine ibex RADseq dataset 14 loci were identified as under selection using the double positive approach but no loci were triple positives. Based on the simulations, a proportion of <0.04 of loci identified by the double positive approach are likely to be true positives. This extremely low proportion indicates that these putatively selected loci should be viewed with extreme caution because many are likely to be false positive loci. Consequently, these loci were not explored further (as in, Shultz *et al.,* 2016). Interestingly, the significant environmental correlations observed with the loci putatively under selection in the Alpine ibex were related to environmental variables known to have recruitment effects and to vary dramatically across the reintroduced range. Despite biologically realistic explanations, the expected high rates of false positives prevent us from making any confident conclusions about local adaptation in the Alpine ibex at this time. Furthermore, the size and nature of this species make the functional validation that was used in *Peromyscus spp.* impossible (Poh *et al.,* 2014). Though it is likely some adaptation may be occurring in Alpine ibex, these candidate outliers and those found in other bottlenecked species, must be confirmed when more accurate selection methods for bottleneck population are identified in the future. Future studies should focus on selection detection methods less reliant on *Fst* (e.g. time series approach, Brüniche-Olsen *et al*., 2016), and explore if sufficient power can be gained by more densely sampling the genome with Whole Genome Sequencing. For studies interested in examining multiple naturally bottlenecked populations (i.e. not reintroduced species) exploiting museum and collection specimens may help circumvent major genetic drift driven false positives by offering pre-bottleneck allele frequencies.

### Simulated data and selection detection accuracy

Alpine ibex have experienced several profound and serial population bottlenecks. Given this extreme history, genome-wide drift effects are highly likely and a high false positive rate was expected for selection detection methods applied to this data (Kimura 1955a; Kimura 1955b; Lotterhos and Whitlock, 2014). The simulations of the Alpine ibex demography confirmed this, revealing an expected false positive rate of up to 0.03 and a false discovery rate often exceeding 0.99 of all outliers. This accuracy was considerably less than that found for non-bottlenecked populations and for scans where a single population is bottlenecked (e.g. 0.1 false positive rate, Foll and Gaggiotti, 2008). However, the low accuracy is similar to studies where more ancient bottlenecks were simulated (e.g. 0.03-0.41 false positive rate, Poh *et al.,* 2014; 0.05-0.30, Shultz *et al.,* 2016). Importantly, increasing stringency to a double or triple positive approach did improve the false positive rate in the Alpine ibex data. This suggests that the double or triple overlap approaches may offer some improved power in bottlenecked populations, and their accuracy should be assessed for more simple bottleneck histories. However, this approach increases the already high risk of being too stringent and removing all loci under selection (high false negative rate), which must also be taken in to account when applying this method.

A low true positive rate was identified for all simulated loci under selection. To generate a biologically realistic trait, majority of loci simulated were of small or moderate effect and it has been previously demonstrated that many selection detection methods struggle to identify such loci, regardless of demographic history (e.g. Biswas and Akey, 2006; Kalsson and Moen, 2010; Narum and Hess, 2011; Kemper *et al*., 2014; Lotterhos and Whitlock, 2015). This is particularly pronounced for loci contributing to polygenic traits such as ours (Kemper *et al.*, 2014; Berg and Coop, 2014). However, in this study, loci under comparable selection coefficients were identified much less frequently than expected based on previous studies. Specifically in our study, loci with a selection coefficient below 0.012 were rarely identified by the double or triple positive method. However, Lotterhos and Whitlock (2015) found a true positive rate of at least 0.11 for loci under a weaker selection coefficient of 0.005, with two or more selection detection methods. Our true positive rate for loci of the largest effect was also lower than seen previously, for example for the Bayenv GEA we found a 0.04 true positive rate, while previous studies have found 0.58-1 across multiple demographic scenarios (Coop *et al.,* 2010; De Mita *et al.,* 2013; Lotterhos and Whitlock; 2015).

The lower accuracy found here is likely driven by a combination of factors, including the intrinsic characteristics of bottlenecked populations. Specifically, the swamping of true positives with drift-driven false positives (which will increase the false discovery and false positive rate), as well as the lower effective population size of a bottlenecked species. A lower effective population size will reduce the efficacy of selection (Frankham *et al.,* 2010). This in turn limits detectable signals of selection. Though 17 thousand Alpine ibex are now present in the Alps, population connectivity is low and contemporary population sizes are often in the hundreds. Effective population sizes range from ~900 to as low as 20 (Biebach and Keller, 2009). While the strength of selection at loci with an allelic value of 0.1 or 0.8 (*s*>0.02) was sufficient to theoretically elicit a response even in the smallest simulated populations (*s*>1/2Ne, Frankham *et al.,* 2010), loci of the smallest effect will not overpower drift unless the effective population size exceeds 125 individuals and the census size of three of our simulated populations fell below this threshold. The reduced efficacy of selection in our smallest populations must disrupt signals of selection at loci under weak selection, and contribute to the low true positive rate observed for these loci. In addition, loci under stronger selection were more often at extreme allele frequencies after the burn-in (i.e. preceding any bottleneck) and their rare alleles were easily lost during the bottlenecks or during the shifts in selection pressures. Many of these loci had to be subsequently excluded from selection scans due to their fixation across all populations, exacerbating our difficulty in identifying selection. These issues are likely common to selection scans on bottlenecked species where selection is long acting (i.e. continuous before and during a bottleneck). Accordingly, true positive rate is similar to that found in other bottlenecked species (e.g. Poh *et al.,* 2014). This result does suggest that greater success may be had when looking for signals of post-bottleneck adaptation, for example when scanning for rapid post-reintroduction adaptation to a novel environmental variable or adaptation to a new disease. Greater success may also be had by using pre-bottleneck samples for SNP ascertainment and implementing temporal selection detection methods.

### Conclusions

Overall, for populations like the Alpine ibex with a history of extreme population bottlenecks (and notably, serial founding events as well as complex reintroductions) the selection detection methods explored here have a considerably reduced accuracy relative to other demographic histories. Based on these results, loci identified as under selection in similar bottlenecked populations using GEA or *Fst* outlier methods should be viewed with caution, particularly those based on single selection detection methods. Unfortunately for bottlenecked species, the high false positive rate is also coupled with a high false negative rate. Therefore, if selective responses are not identified in bottlenecked populations this cannot be considered evidence for an absence of selection pressures or an absence of local adaptation. This unfortunate lack of power, is highly problematic for effective adaptive population management. However, the costs of falsely concluding two populations as separate ESUs based on erroneous outliers could be high. The criteria and approach outlined here may offer other studies on bottlenecked species an approach and baseline on which to gauge their confidence in any outliers identified and adjust management plans accordingly. In the future, selection detection methods less reliant on *Fst,* such as inter-species comparisons or those exploiting temporal samples (Brüniche-Olsen *et al*., 2016), as well as use of more dense marker data, should be explored across more bottlenecked scenarios. Despite the high false positive rate expected, it is important to see if these approaches offer greater power and if they can better facilitate conservation management.

## Acknowledgements

This project was funded by a University of Zurich’s Research Priority Program “Evolution in Action” grant to Lukas F Keller and Andreas Wagner. We would like thank them both for their support throughout this project. We would also like to thank the European Science Foundation for funding a Short Visit Grant to DML and TG to establish the collaboration. We would like to thank the ETH’s Genetic Diversity Center and BSSE for their help generating the RADseq data. We would also like to thank Glauco Camenisch, Kasia Sluzek and Iris Biebach for their help during the project.

## Data accessibility

Read data can be viewed on the short-read archive, ncbi project number PRJNA422727: https://www.ncbi.nlm.nih.gov/bioproject/PRJNA422727. The demultiplexing file is accessible at http://datadryad.org/review?doi=doi:10.5061/dryad.8vm8d.

## Author contributions

DML performed the selection detection analysis, simulations and wrote the manuscript

TG supported the selection detection analysis and commented on the manuscript.

CG supported the sequence data generation, commented on the manuscript and simulations.

FG supported writing the simulation scripts and designing the genetic architecture of the QTL traits.

