## Supplementary Material for "Disentangling adaptation from drift in bottlenecked and reintroduced populations of Alpine ibex"

Running title: Outlier scan accuracy in bottlenecked populations

Leigh, D.M.,* Lischer, H.E.L., Guillaume, F., Grossen, C., Günther, T.

#### Supplementary Material

**Section S1:**

**Simulations of neutral loci and loci under selection:**

Nemo (version 2.3.51; Guillaume and Rougemont, 2006) was used to create forward simulations where individuals follow a user-determined life cycle of reproduction, migration and breeding partner numbers. Genotypes were generated for each individual and were sampled at the end of the simulations. 26 populations were simulated, representing the 23 focal Swiss Alpine ibex populations, the Gran Paradiso (source population) and two additional captive populations used as founder sources. To reach mutation-drift equilibrium and enable a build up of linkage disequilibrium, an initial run of 10,000 generations (a 'burn-in') was conducted for the simulated population representing the Gran Paradiso remnant ibex population. After reaching equilibrium, a bottleneck was applied to mirror the global decline in Alpine ibex. The Gran Paradiso population was reduced to 80 individuals and then allowed to regrow over 7 generations, after which reintroductions began. The time from the first reintroduction in 1906 to 2015 was divided into discrete generations of eight years. Eight years corresponds to the generation time of Alpine ibex (Stuwe and Grodinsky, 1987). All individuals that were reintroduced in each eight-year period were represented as one migration event in the corresponding Nemo generation. The period from 1906 to 2015 corresponds to 13.6 generations. Accordingly, to ensure that all translocations were included in the simulations, 14 generations of reintroductions were simulated. The last five-year period corresponds to one full generation. In addition, a final generation was allowed after the 14 generations of reintroductions, so that the last reintroduced individuals could reproduce before sampling. No migration outside of recorded translocations has been detected genetically (Aeschbacher *et al.,* 2013). Therefore, no additional migrations were allowed between simulated populations. Five minor contributing source populations in were ignored in the simulations, as this would have created further complexity arising from the need to simulate their founders in turn. The effect of ignoring these populations should be negligible because genetically very similar populations were available to use as source population for the simulated translocations. Only very minor reintroductions (fewer than six individuals) from these non-simulated populations were ignored.

Migration events within Nemo were determined from migration rates. Specifically, we determined the proportion of the source population individuals that migrate. Consequently, for any simulated movement of individuals that represented a translocation, the proportion of individuals that migrated was monitored to ensure this was as similar as possible to the size of the translocation event. One source population was founded a generation early to ensure migration from that population could occur at the correct generation. Populations were permitted to grow to a pre-determined maximum size, after which individuals were randomly removed to keep the population size constant. The maximum size was the largest census size observed between the start of recorded population sizes and the year 2006. 2006 was chosen as it corresponded to the start of the final generation within simulations and the start of the genetic sampling. However, it should be noted that Alpine ibex populations often attained their maximum population size well before 2006 in the wild. The maximum Alpine ibex population size used excluded individuals less than one-year-old where possible because Nemo only regulates adult numbers (Guillaume and Rougemont, 2006). No census data was available for three populations, consequently their maximum size was set to 319 individuals, the mean maximum population size across Swiss populations with census records. All individuals were given a mean lifetime fecundity of 10 offspring to prevent early population extinction in the reintroduction phase of the simulations. Reproductive success followed a Poisson distribution and males were randomly chosen for each female. In the wild, Alpine ibex have a strong skew in reproductive successes to a handful of older males in a population (Willisch *et al.,* 2012). Unfortunately, at the time of the analysis, a similar option was not available in Nemo, which is why randomly chosen males were used instead.

Ten replicate 'burn-ins' were conducted for each genetic architecture (neutral only, 120 selected loci or 30 selected loci). These populations were then used as starting points for 10 bottleneck replicate simulations for each architecture. For each simulation the evolution of 60,000 SNPs on 30 chromosomes of equal size was simulated. 60,000 SNPs ensured that enough SNPs were polymorphic at the end of the simulations to be randomly subsampled to match the empirical RADseq data set. SNPs were evenly distributed and recombined at a rate of $5x10^{-5}$. This created a genome similar in size and structure to that of the goat (Guillaume and Rougemont, 2006; Bickhart *et al.,* 2017). All SNPs were initially monomorphic and variation arose at a mutation rate ($\mu$) equal to $1 \times 10^{-6}$ per locus and generation. After the burn-in no new mutational input was allowed.

To simulate selection, 30 or 120 additional SNPs were placed individually on each chromosome, each of these contributed additively to a single trait. Allelic values of 0.1, 0.8, 0.4, 0.2 and 0.1 were simulated. During the burn-in a local trait optimum of zero was used to maintain alleles under selection that had both positive and negatives contributions on the trait value. To maintain polymorphisms at the loci under selection a higher mutation rate of 1 $x10^{-4}$ was used. For the architecture with 120 loci under selection, a genetic map resolution of 0.05 was used and loci were at position, 10, 6670, 13330 or 19990 on the genetic map. For the architecture with 30 loci under selection the same resolution and positions were available but only one was used per chromosome. Population optimum trait values for reintroduced populations were set to reflect the patterns seen in natural environmental conditions. Specifically, the relative recorded difference in the reintroduced populations snow conditions determined whether it was set to -2, +2 or remained at 0. Simulated optima are shown in Table S1.

For selection detection analyses a maximum of ten individuals from 23 populations were selected. Fixed SNPs were excluded from this analysis. Additionally, the three populations that were not sampled genetically but were simulated as founder sources were also excluded from the search for signals of positive selection in the simulated data. All genotypes were taken from the final generation of the simulations.

**Section S2:**

**Weather Data:**

To include a measure of the environmental conditions a population experienced, the environmental conditions of the closest meteorological weather station were averaged across years. If weather records began before a population was founded, data were averaged from the year after the first translocation event until October 2015, the end of population sampling. For populations where weather records began after the first translocation event, all data were used. Each environmental variable included was divided into three seasons: winter (November-April), summer (May-October), and late winter (February to April) (Buzzuto*, pers.com*). Late winter was included as it captures the period with highest snow depths. The environmental variables included in the analyses were: daily mean, maximum and minimum air temperature 2 meters above ground (°C), daily precipitation (mm), daily new snow accumulation (cm), and total snow height (cm) at 5:40 am (MeteoSwiss, Switzerland). Mean values were calculated separately for each season and used in Bayenv 2.0 and Baypass2.1. For the simulated data sets the mean value of each environmental variable and season corresponding to the simulated population were used in the GEA approaches.

**Section S3:**

**RADseq bioinformatics steps:**

RADseq data generation and initial processing is detailed in Leigh *et al.,* (2018) and Grossen *et al.,* (2017) (read data NCBI number PRJNA422727). Following the recommendations in Leigh *et al.,* (2018) variant nucleotides were called using GATK's HaplotypeCaller (version 3.4-46-gbc02625, Poplin *et al.,* 2017). SNPs only polymorphic between the Alpine ibex and the goat genome were removed before filtering using custom scripts (available on dryad, https://doi.org/10.5061/dryad.8vm8d). SNPs were then filtered with GATK's 'VariantFiltration.' To remove poor quality variants, SNPs were removed if the site quality to depth (QD) value was less 2, if the FisherStrand (FS) bias value was greater than 60, if site mapping quality was less than 40, if the MappingQualityRankSumTest (MQRankSum) was less then -4.0, if the

ReadPositionRankSum was less than -4.0 and if the StrandsOddsRatio (SOR) was greater than 3. To avoid biases arising from different library lengths at the read-ends (see Leigh et al., 2018), SNPs were removed at read ends SNPs with a RPRS value of less than -4 or greater than 4 were removed. Vcftools was used to remove SNPs with a genotyping quality of less than 20 and a depth of less than 8, to remove remaining poor quality variants. To remove uninformative SNP sites with over 50% of genotypes missing were excluded. To ensure paralogs and duplicate regions were not present (Li, 2014), SNPs with a mean maximum read depth across all individuals greater than twice the mean depth (>31) and those with a heterozygosity of greater than 0.6 were excluded. Finally, SNPs with more or less than two alleles were removed, because the selection detection programs could not accept non-biallelic loci. Two individuals were excluded after filtering due to abnormal heterozygosity values.

***Table S1:*** *The simulated optima of the populations included in the simulations. Positive and negative values were based on the relative difference in snow depth between each population and the Gran Paradiso population. An optimum of zero was used for the burn-in.*

| Population | Simulated optimum |
|---|---|
| Gran Paradiso | 0 |
| Wilderness Park Peter and Paul | -2 |
| Wilderness Park Interlaken Harder | -2 |

| | |
|---|---|
| Graue Hoerner | -2 |
| Albris | 0 |
| Brienzer-Rothorn | 0 |
| Schwarmoench | -2 |
| Wetterhorn | 0 |
| Mont Pleureur | -2 |
| Justistal | 0 |
| Gross Lohner | -2 |
| Alpstein | 2 |
| Val Bever | -2 |
| Crap da Flem | -2 |
| Flueela | 0 |
| Wittenberg | 0 |
| Arolla | -2 |
| Bire-Oeschinen | -2 |
| Creux du Van | 0 |
| Pilatus | 0 |
| Fluebrig | -2 |
| Weisshorn | -2 |
| Oberbauenstock | -2 |
| Falknis | 0 |
| Tanay | -2 |
| Churfirsten | 2 |

***Table S2:*** *Weather stations used for each population. Data included in the analyses are an average of each season over the years since each population's founding or when records began. Stations were chosen based on proximity and similarity in conditions.*

| **Colony** | **First Translocation** | **Weather Time Series Start** | **Station** | **Station Elevation (m)** |
|---|---|---|---|---|
| Bire-Oeschinen | 1961 | 1962 | ABO | 1320 |
| Gross Lohner | 1952 | 1953 | ABO | 1320 |
| Schwarmoench | 1924 | 1931 | ABO | 1320 |
| Tanay | 1977 | 1978 | AIG | 381 |
| Albris | 1920 | 1931 | BEH | 2307 |
| Crap da Flem | 1958 | 1959 | ELM | 965 |
| Fluebrig | 1962 | 1963 | ELM | 965 |
| Graue Hoerner | 1911 | 1912 | ELM | 965 |
| Oberbauenstock | 1969 | 1970 | ENG | 1037 |
| Arolla | 1960 | 1978 | EVO | 1825 |
| Creux du Van | 1961 | 1978 | FRE | 1205 |
| Wetterhorn | 1926 | 1931 | GRH | 1980 |
| Mont Pleureur | 1928 | 1928 | GSB | 2472 |
| Wittenberg | 1958 | 1978 | MLS | 1972 |
| Brienzer-Rothorn | 1921 | 1978 | PIL | 2106 |
| Justistal | 1949 | 1978 | PIL | 2106 |

| | | | | |
|---|---|---|---|---|
| Pilatus | 1961 | 1978 | PIL | 2106 |
| Alpstein | 1955 | 1956 | SAE | 2502 |
| Churfirsten | 1984 | 1985 | SAE | 2502 |
| Falknis | 1970 | 1971 | WFJ | 2690 |
| Flueela | 1958 | 1959 | WFJ | 2690 |
| Val Bever | 1957 | 1978 | SAM | 1705 |
| Weisshorn | 1962 | 1962 | ZER | 1638 |

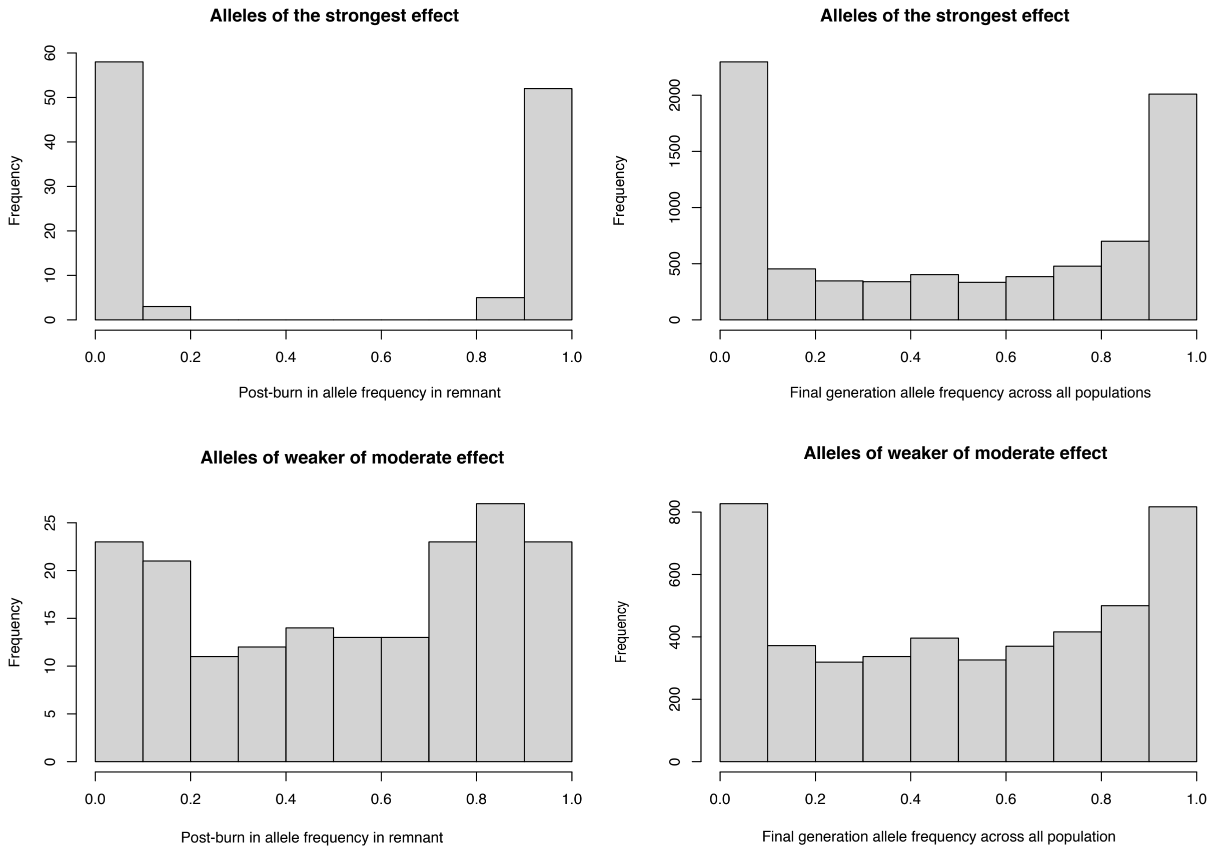


***Figure S1:*** *Allele frequencies of selection loci from the architecture with 30 loci under selection. Shown are loci that remain polymorphic after the burn-in within the remnant population (left panels) and their subsequent frequencies across all populations (right panels) in the final simulated generation. Top panels are loci under strong selection with an allelic value of 0.08 or 0.1, bottom panels are loci under weaker selection with an allelic value of 0.04,0.02 or 0.01.*

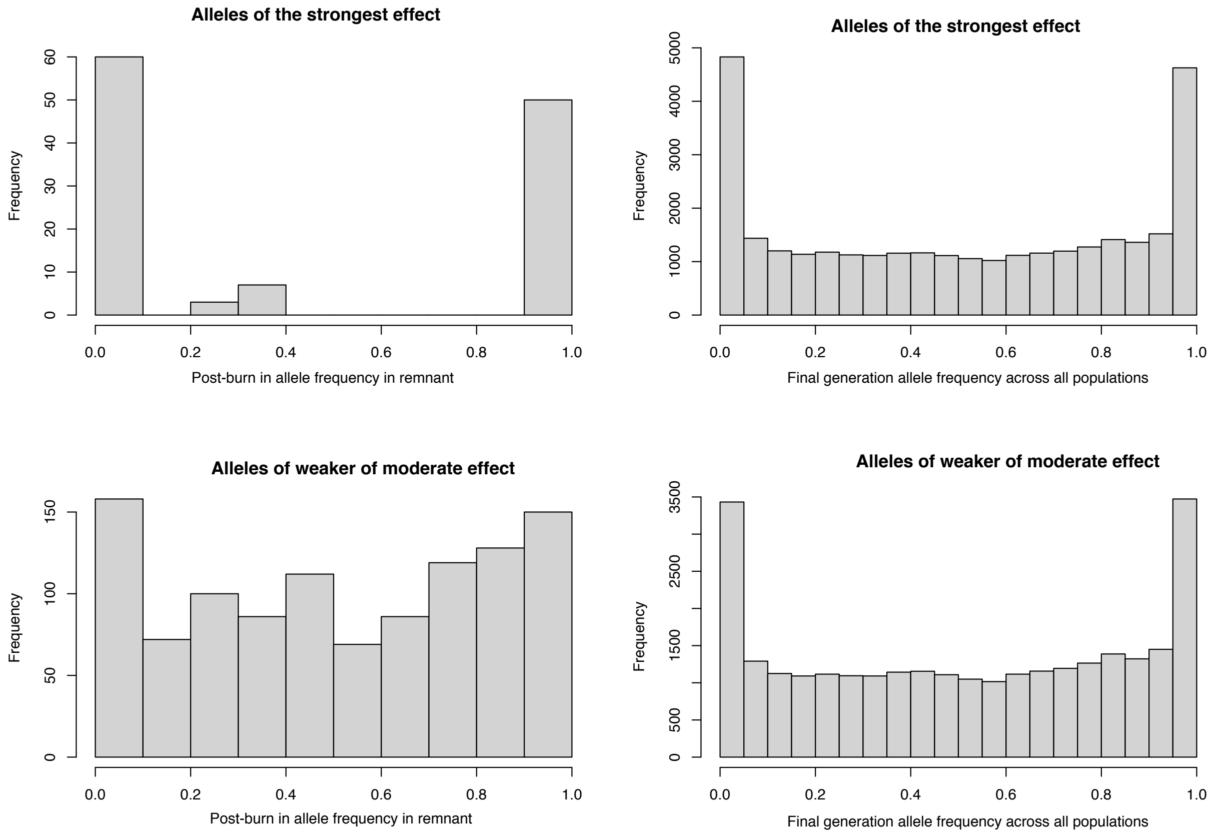


***Figure S2:*** *Allele frequencies of selection loci from the architecture with 120 loci under selection. Shown are loci that remain polymorphic after the burn-in within the remnant population (left panels) and their subsequent frequencies across all populations (right panels) in the final simulated generation. Top panels are loci under strong selection with an allelic value of 0.08 or 0.1, bottom panels are loci under weaker selection with an allelic value of 0.04,0.02 or 0.01.*

### Supplementary Material References

Aeschbacher, S., Futschik, A., & Beaumont, M.A. (2013). Approximate Bayesian computation for modular inference problems with many parameters: the example of migration rates. Molecular ecology, 22(4), 987–1002. doi:10.1111/mec.12165

Ahn, S.J., Costa, J., & Emanuel, J.R. (1996). PicoGreen quantitation of DNA: effective evaluation of samples pre- or post-PCR. Nucleic Acids Res. 24:2623–2625.

Bickhart, D.M., Rosen, B.J., Koren, S., Sayre, B.L., Hastie, A.R.,…Smith, T.P.L. (2016) Single-molecule sequencing and chromatin conformation capture enable de novo reference assembly of the domestic goat genome. Nature Genetics. 49:643-650 doi:10.1038/ng.3802

Bolger, A.M., Lohse, M., & Usadel, B. (2014). Trimmomatic: a flexible trimmer for Illumina sequence data. Bioinformatics. 30:2114–2120. 10.1093/bioinformatics/btu170

Broad (2016) Picard. Retrieved from http://broadinstitute.github.io/picard.

Danecek, P., Auton, A., Abecasis, G., Albers, C.A., Banks, E., …Wang, J. (2011). The variant call format and VCFtools. Bioinformatics 27:2156–8. 10.1093/bioinformatics/btr330

Dong, Y., Xie, M., Jiang, Y., Xiao, N., Du, X., Zhang, W., ... Wang, W. (2013). Sequencing and automated whole-genome optical mapping of the genome of a domestic goat (Capra hircus). Nature Biotechnology. 31:135–41. doi:10.1038/nbt.2478

Etter, P.D., Bassham, S., Hohenlohe, P.A., Johnson, E.A., & Cresko, W.A. (2011). Molecular Methods for Evolutionary Genetics. In: Orgogozo V, Rockman MV, editors. Vol. 772. Totowa, NJ: Humana Press. p. 157–178.

Grossen, C., Keller, L., Biebach, I., & Croll, D. (2014). Introgression from domestic goat generated variation at the major histocompatibility complex of Alpine ibex. PLoS Geneteics. 10(6),1-19, 10:e1004438

Grossen, C., Biebach, I., Angelone-Alasaad, S., Keller, L. F., & Croll, D. (2017). Population genomics analyses of European ibex species show lower diversity and higher inbreeding in reintroduced populations. Evolutionary Applications.11:123–139

Guillaume, F., & Rougemont, J. (2006). Nemo: an evolutionary and population genetics programming framework. Bioinformatics. 22:2556–7 10.1093/bioinformatics/btl415

Langmead, B., & Salzberg, S.L. (2012). Fast gapped-read alignment with Bowtie 2. Nature Methods 9:357–9.

Leigh, D. M., H. E. L. Lischer, C. Grossen, and L. F. Keller. 'Batch Effects in a Multiyear Sequencing Study: False Biological Trends Due to Changes in Read Lengths'. Molecular Ecology Resources 18, no. 4 (2018): 778–88. https://doi.org/10.1111/1755-0998.12779.

Li, H. 2014. Toward better understanding of artefacts in variant calling from high-coverage samples. Bioinformatics 30:2843–51. 10.1093/bioinformatics/btu356

Pearson, W.R., Wood, T., Zhang, Z., & Miller, W. (1997). Comparison of DNA Sequences with Protein Sequences. Genomics. 36:24–36. dx.doi.org/10.1006/geno.1997.4995

Poplin, R., Ruano-Rubio, V., DePristo,M.A., Fennell, T.J., Carneiro, M.O., Van der Auwera, G.A., … Banks, E. (2017) Scaling accurate genetic variant discovery to tens of thousands of samples *bioRxiv*

Stuwe, M., & Grodinsky, C. (1987). Reproductive biology of captive Alpine ibex (*Capra i. ibex*). Zoo Biology. 6,331–339. 10.1002/zoo.1430060407

Willisch, C., I. Biebach, U. Koller, T. Bucher, N. Marreros,…Neuhaus, P. (2012) Male reproductive pattern in a polygynous ungulate with a slow life-history: the role of age, social status and alternative mating tactics. Evol. Ecol. 26,187–206
